# Mosaic Epigenetic Dysregulation of Ectodermal Cells in Autism Spectrum Disorder

**DOI:** 10.1101/004853

**Authors:** Esther R. Berko, Masako Suzuki, Faygel Beren, Christophe Lemetre, Christine M. Alaimo, R. Brent Calder, Karen Ballaban-Gil, Batya Gounder, Kaylee Kampf, Jill Kirschen, Shahina B. Maqbool, Zeineen Momin, David M. Reynolds, Natalie Russo, Lisa Shulman, Edyta Stasiek, Jessica Tozour, Maria Valicenti-McDermott, Shenglong Wang, Brett S. Abrahams, Joseph Hargitai, Dov Inbar, Zhengdong Zhang, Joseph D. Buxbaum, Sophie Molholm, John J. Foxe, Robert W. Marion, Adam Auton, John M. Greally

## Abstract

DNA mutational events are increasingly being identified in autism spectrum disorder (ASD), but the potential additional role of dysregulation of the epigenome in the pathogenesis of the condition remains unclear. The epigenome is of interest as a possible mediator of environmental effects during development, encoding a cellular memory reflected by altered function of progeny cells. Advanced maternal age (AMA) is associated with an increased risk of having a child with ASD for reasons that are not understood. To explore whether AMA involves covert aneuploidy or epigenetic dysregulation leading to ASD in the offspring, we tested an homogeneous ectodermal cell type from 47 individuals with ASD compared with 48 typically developing (TD) controls born to mothers of ≥35 years, using a quantitative genome-wide DNA methylation assay. We show that DNA methylation patterns are dysregulated in ectodermal cells in these individuals, having accounted for confounding effects due to subject age, sex and ancestral haplotype. We did not find mosaic aneuploidy or copy number variability to occur at differentially-methylated regions in these subjects. Of note, the loci with distinctive DNA methylation were found at genes expressed in the brain and encoding protein products significantly enriched for interactions with those produced by known ASD-causing genes, representing a perturbation by epigenomic dysregulation of the same networks compromised by DNA mutational mechanisms. The results indicate the presence of a mosaic subpopulation of epigenetically-dysregulated, ectodermally-derived cells in subjects with ASD. The epigenetic dysregulation observed in these ASD subjects born to older mothers may be associated with aging parental gametes, environmental influences during embryogenesis or could be the consequence of mutations of the chromatin regulatory genes increasingly implicated in ASD. The results indicate that epigenetic dysregulatory mechanisms may complement and interact with DNA mutations in the pathogenesis of the disorder.

**AUTHOR SUMMARY:** Older mothers have a higher than expected risk of having a child with an autism spectrum disorder (ASD). The reason for this increased risk is unknown. The eggs of older mothers are more prone to abnormalities of chromosome numbers, suggesting this as one possible mechanism of the increased ASD risk. Age is also associated with a loss of control of epigenetic regulatory patterns that govern gene expression, indicating a second potential mechanism. To test both possibilities, we sampled cells from the same developmental origin as the brain, and performed genome-wide tests looking for unusual chromosome numbers and DNA methylation patterns. The studies were performed on individuals with ASD and typically developing controls, all born to mothers at least 35 years of age at the time of birth. We found the cells from individuals with ASD to have changes in DNA methylation at a number of loci, especially near genes encoding proteins known to interact with those already implicated in ASD. We conclude that epigenetic dysregulation occurring in gametes or early embryonic life may be one of the contributors to the development of ASD.

## INTRODUCTION

Progress in understanding the genetic basis of ASD has been substantial in recent years, with the development of microarray technologies allowing the identification of copy number variants associated with the disorder [1] and massively-parallel sequencing focused on protein-coding exons allowing insights into smaller mutational events disrupting gene function [2,3]. The emerging picture is of rare rather than common genetic variants mediating most of the risk [4]. Less progress has been made in understanding the mechanism by which environmental factors influence the risk of ASD [5]. Epigenetic mediation of such environmental influences has been proposed [6], but a clear association between the phenotype and epigenetic dysregulation has proven elusive in genome-wide studies [7]. Three recent studies have lent support to an association of epigenetic dysregulation with ASD. One study found indications of distinctive chromatin features in the ASD brain [8], another tested peripheral blood leukocytes and found DNA methylation differences to characterize monozygotic twins affected by ASD compared with their unaffected twin [9], while a third study tested brains from 19 subjects with ASD and 21 controls, also finding differential DNA methylation [10].

Interestingly, chromatin regulatory genes have been described to be significantly enriched as targets of mutational events [11], suggesting that epigenomic dysregulation secondary to mutational events may mediate pathophysiological dysfunction in some individuals with ASD. We were also interested in the parental age effect in ASD, which for advanced paternal age appears to be substantially attributable to mutational events in the male germline [12,13]. The mechanism of the independent maternal age effect on ASD prevalence [14] has substantially less evidence for such underlying genetic events and remains mechanistically unclear. As advanced maternal age (AMA) has long been associated with increased rates of chromosomal non-disjunction [15], while cellular aging in general is increasingly recognized to involve epigenetic dysregulation [16], we therefore evaluated both of these mechanisms as potential contributors to the increased risk of ASD in a cohort of individuals born to older mothers.

A major component of the current study was the minimization of the potential confounding effects now increasingly recognized to affect epigenome-wide association studies (EWAS) [17,18]. We therefore ensured that we reduced effects due to cell type and subpopulation heterogeneity, chromosomal aneuploidy, copy number variability, genetic polymorphism, age, sex and technical influences, and defined differential DNA methylation using advanced analytical approaches. Our intent was to perform a study representing the best of current practices, maximizing our chances of identifying epigenetic regulatory patterns associated with a complex disorder like ASD.

## RESULTS

### Minimizing the effects of confounding influences on the DNA methylation data

The conclusions of our study are based on Co-methylation Network Analysis approaches described later, but we also included more mainstream analytical techniques to demonstrate that these also yield information about differential DNA methylation characterizing the ASD group. As a first pass analytical approach, we performed on our preprocessed dataset the type of non-parametric F-testing typically performed for Illumina 450K DNA methylation microarrays (IMA [19]), identifying 3,560 differentially methylated CGs (p<0.001) discriminating ASD and TD individuals. We were, however, concerned that this kind of analysis would be subject to confounding influences and prove to be misleading. Such confounding influences include subject age [16,20,21], gender [20,22], and genetic sequence polymorphism at the tested site or acting *in cis* [23-25]. We therefore sought to understand the contribution of these covariates on our dataset. To accomplish this, we adapted an approach used in conjunction with principal components analysis [26]. We assessed all known technical and biological covariates, and then further determined ancestral haplotype of each individual by analyzing genotype data from cohort subjects and applying the *HAPMIX* algorithm (Figure 1, **Supplemental Table S3**). We then performed linear regression of each principal component on each covariate, to determine significant correlations (Figure 2). We found that inter-array differences, reflected by bisulphite conversion and DNA loading controls on the microarray, accounted for the majority of confounding variation between samples, even after applying stringent preprocessing algorithms. We also found that subject age and ancestry contributed significantly to DNA methylation variation.

**Figure 1:**
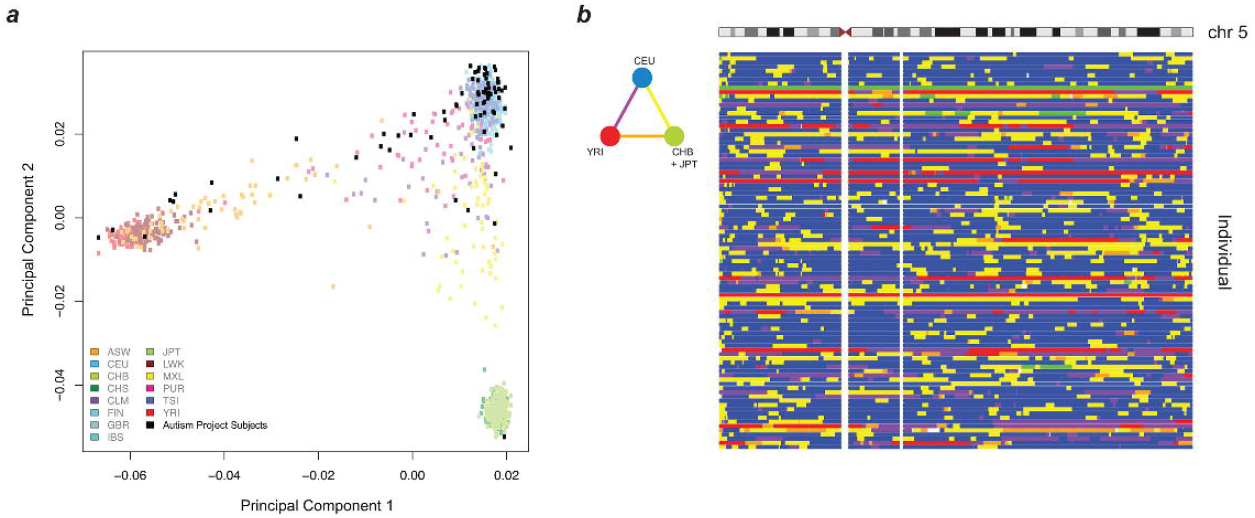
Principal components analysis (PCA) of ASD project genotypes with 1000 Genomes data and local ancestry deconvolution across chromosome 5. A. **Autism Project Subjects**: We show 78 out of 95 non-founders in the cohort, after removing siblings for the PCA and keeping only one member per family. **ASW**: Americans of African Ancestry in SW USA; **CEU**: Utah Residents (CEPH) with Northern and West European ancestry; **CHB**: Han Chinese in Beijing, China; **CHS**: Southern Han Chinese; **CLM**: Columbians from Medellin, Colombia; **FIN**: Finnish in Finland; **GBR**: British in England and Scotland; **IBS**: Iberian population in Spain; **JPT**: Japanese in Tokyo, Japan; **LWK**: Luhya in Webuye, Kenya; **MXL**: Mexican Ancestry from Los Angekes USA; **PUR**: Puerto Ricans from Puerto Rico; **TSI**: Toscani in Italia; **YRI**: Yoruba in Ibadan, Nigeria.
B. The triangular color key denotes the color scheme used for genotype. **Blue**: Homozygous CEU; **Green**: Homozygous CHB + JPT; **Red**: Homozygous YRI; **Yellow**: Heterozygous CEU/CHB + JPT; **Orange**: Heterozygous CHB + JPT/YRI; **Purple**: Heterozygous YRI/CEU.

**Figure 2:**
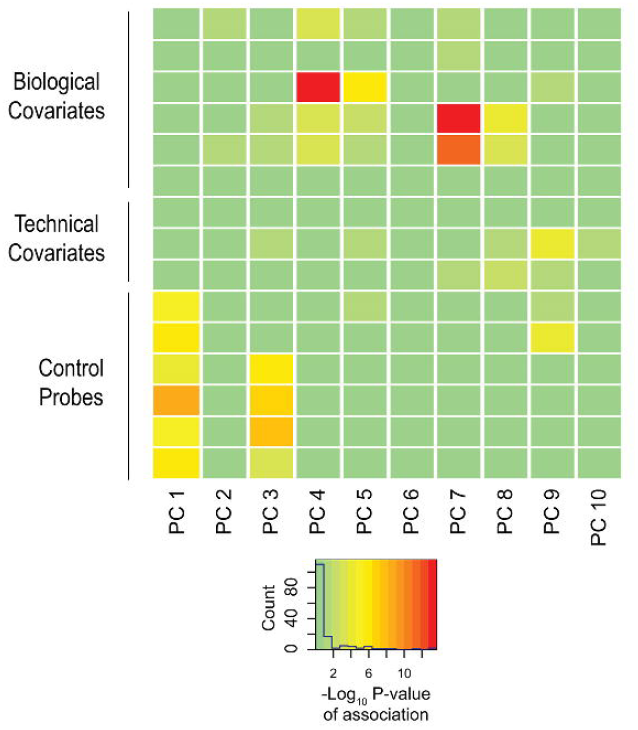
Biological and technical confounders contribute to methylation value variation. The heat map displays the –log_10_ p-values of the linear regressions of the top ten principal components onto each known covariate. The color key shows corresponding numeric values, with red indicating increased significance. The majority of variation is accounted for by experimental influences, with age and ancestry also contributing significantly to variation.

To strengthen the quality of our dataset further, we used our genotype data to explore the presence of detectable mosaic aneuploidy and copy number variation in the subjects. We applied the Mosaic Alteration Detection (*MAD*) algorithm to test for mosaicism for chromosome aneuploidy [27], finding no evidence for such an event in our cases or controls. We were therefore able to focus on the analysis of DNA methylation in the same samples knowing that the results would not be confounded by AMA-induced aneuploidy.

### Identifying Differentially Methylated Regions (DMRs)

To identify DNA methylation changes specific to the ASD subjects, we used two approaches, the identification of differentially methylated regions (DMRs) using the bump-hunting approach [28], and, as we describe in the next section, Co-methylation Network Analysis. DMR identification has the advantage of being conceptually easy to understand and lends itself to single locus verification studies, as it focuses on the definition of discrete loci with DNA methylation differences. Instead of interrogating individual sites, bump-hunting combines probe information over short regions of hundreds of basepairs, and therefore minimizes single locus false positives that can occur due to polymorphism at individual CGs. Importantly, unlike approaches like IMA [19], bump-hunting considers the many potential sources of confounding variation in epigenomic analysis and includes analytical approaches to account for and remove unwanted covariates. Bump-hunting was previously developed for tiling microarray data [28]; our use of the algorithm for the Infinium 450K microarray platform represents an extension of the approach that has proven successful in another study of ASD [10]. Applying the default 300bp window [28], this allows ∼27% of the probes on the Infinium 450K microarray to be informative.

Since we had confirmed effects of age and ancestry on DNA methylation in our subjects, we included them as covariates in the bump-hunting model, estimating ancestry as the percent of European (CEU) and of African (YRI) alleles genome-wide. Removing CGs that overlap known common SNPs from further analysis is a commonly performed measure to reduce artifactual effects of unrecognized sequence variants [29], but is insufficient to account for genotype-driven differences, as recent studies have demonstrated the ability of haplotypes *in cis* with tested CGs to affect their methylation status [20,23,24]. Although we did not observe significant variation due to sex, it was also included as a covariate as a conservative precaution given its previously documented effects [22]. We confirmed that bump-hunting accounted for and removed all of the technical variation by repeating our PCA and association analysis on the bump-hunting output (**Supplemental Figure S4**).

We discovered 15 DMRs at 14 genes distinguishing the ASD and TD samples (**Supplemental Table S4**). As it is known that the incidence of genic copy number variation (CNV) is increased in individuals with ASD [2], we used *CNVision* [30] to identify large CNVs from our genotyping microarray data and to test whether any of the 15 DMRs overlapped CNVs. Three DMRs (one at the *MAPK8IP3* gene and two at the *CYP2E1* gene) were found to overlap CNVs in multiple cases and/or controls. We therefore excluded the individuals with CNVs at those loci and re-ran the DNA methylation bump-hunting analysis. The two candidate DMRs at *CYP2E1* were no longer identified in this re-analysis, indicating that the *CYP2E1* DMR assignation was likely due to the presence of the multiple CNVs.

As an extra layer of stringency, we then re-ran the data preprocessing and analysis for a total of 4 iterations, testing to see which of the remaining 13 loci remained stably predicted as DMRs. We found four (at the *KCNQ5*, *NRG2*, *LOC643802* and *ZG16B* genes) to be predicted in only a subset of the iterative analyses, but the other 9 DMRs to remain stably predicted, generating higher confidence predictions for these 9 loci as a consequence. The instability of prediction of certain loci is likely to be due to *ComBat* generating slightly different outputs every time it is run, requiring this kind of iterative analysis to test for stability of predictions.

The candidate DMRs from the genome-wide analysis were associated with genes, of which many have already been implicated in previous studies with ASD (Table 1). Of the 9 genes defined by the candidate DMRs in buccal epithelial cells, all are expressed in the brain according to the Human Brain Transcriptome (HBT) database [31]. Furthermore, 6 of the 9 genes were ascribed by the HBT to co-expression modules; all 6 belong to 2 modules involved specifically in post-natal synaptic transmission [31]. This includes the *OR2L13* gene, which, unlike other olfactory receptor genes, is not located in the module associated with olfactory function.

**Table 1:**
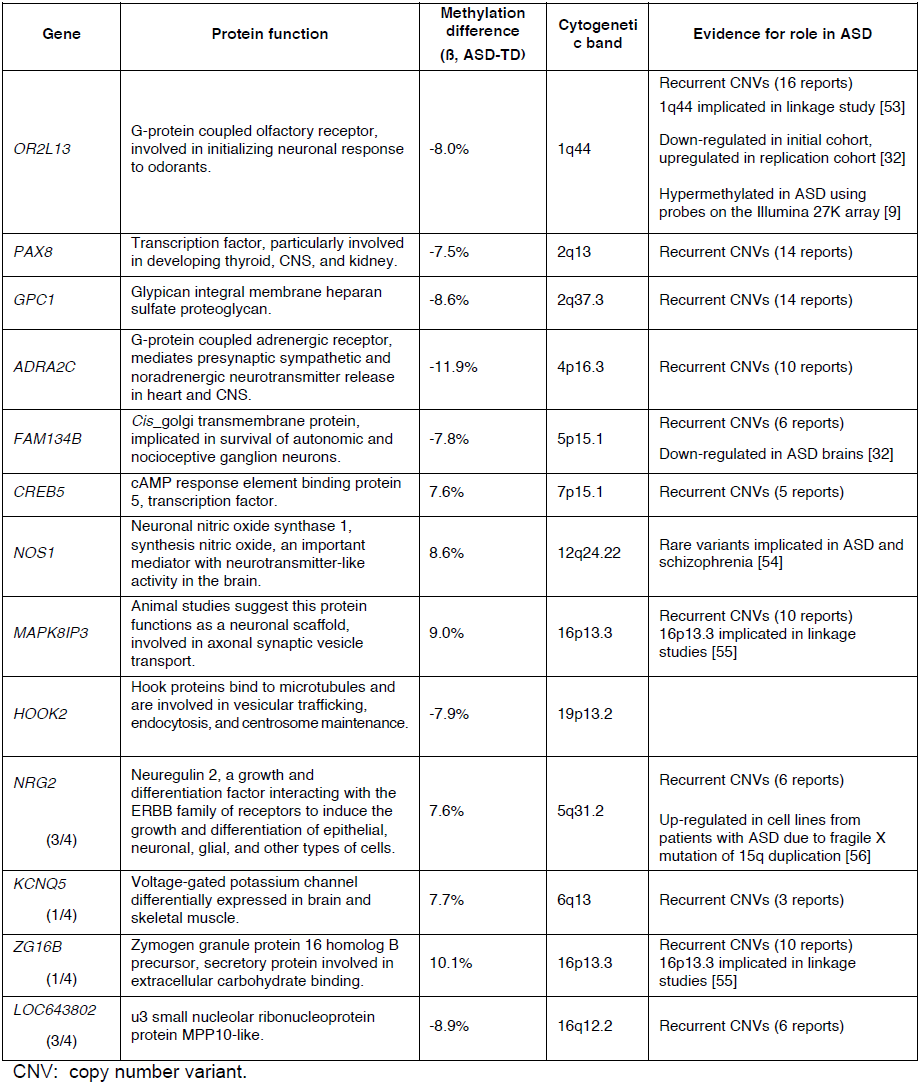
Genes associated with the DMRs identified by bump-hunting.

| Gene | Protein function | Methylation difference (B, ASD-TD) | Cytogenetic band | Evidence for role in ASD |
| --- | --- | --- | --- | --- |
| *OR2L13* | G-protein coupled olfactory receptor, involved in initializing neuronal response to odorants. | -8.0% | 1q44 | Recurrent CNVs (16 reports)<br>1q44 implicated in linkage study [53]<br>Down-regulated in initial cohort, upregulated in replication cohort [32]<br>Hypermethylated in ASD using probes on the Illumina 27K array [9] |
| *PAX8* | Transcription factor, particularly involved in developing thyroid, CNS, and kidney. | -7.5% | 2q13 | Recurrent CNVs (14 reports) |
| *GPC1* | Glypican integral membrane heparan sulfate proteoglycan. | -8.6% | 2q37.3 | Recurrent CNVs (14 reports) |
| *ADRA2C* | G-protein coupled adrenergic receptor, mediates presynaptic sympathetic and noradrenergic neurotransmitter release in heart and CNS. | -11.9% | 4p16.3 | Recurrent CNVs (10 reports) |
| *FAM134B* | *Cis_golgi* transmembrane protein, implicated in survival of autonomic and nociceptive ganglion neurons. | -7.8% | 5p15.1 | Recurrent CNVs (6 reports)<br>Down-regulated in ASD brains [32] |
| *CREB5* | cAMP response element binding protein 5, transcription factor. | 7.6% | 7p15.1 | Recurrent CNVs (5 reports) |
| *NOS1* | Neuronal nitric oxide synthase 1, synthesis nitric oxide, an important mediator with neurotransmitter-like activity in the brain. | 8.6% | 12q24.22 | Rare variants implicated in ASD and schizophrenia [54] |
| *MAPK8IP3* | Animal studies suggest this protein functions as a neuronal scaffold, involved in axonal synaptic vesicle transport. | 9.0% | 16p13.3 | Recurrent CNVs (10 reports)<br>16p13.3 implicated in linkage studies [55] |
| *HOOK2* | Hook proteins bind to microtubules and are involved in vesicular trafficking, endocytosis, and centrosome maintenance. | -7.9% | 19p13.2 |  |
| *NRG2*<br>(3/4) | Neuregulin 2, a growth and differentiation factor interacting with the ERBB family of receptors to induce the growth and differentiation of epithelial, neuronal, glial, and other types of cells. | 7.6% | 5q31.2 | Recurrent CNVs (6 reports)<br>Up-regulated in cell lines from patients with ASD due to fragile X mutation of 15q duplication [56] |
| *KCNQ5*<br>(1/4) | Voltage-gated potassium channel differentially expressed in brain and skeletal muscle. | 7.7% | 6q13 | Recurrent CNVs (3 reports) |
| *ZG16B*<br>(1/4) | Zymogen granule protein 16 homolog B precursor, secretory protein involved in extracellular carbohydrate binding. | 10.1% | 16p13.3 | Recurrent CNVs (10 reports)<br>16p13.3 implicated in linkage studies [55] |
| *LOC643802*<br>(3/4) | u3 small nucleolar ribonucleoprotein protein MPP10-like. | -8.9% | 16q12.2 | Recurrent CNVs (6 reports) |

CNV: copy number variant.

To confirm differential DNA methylation at these candidate DMRs, we used bisulphite PCR, multiplexed robotic library preparation and massively-parallel sequencing averaging >15,000x depth for each locus tested, with >99.99% bisulphite conversion efficiency. We show in Figure 3 the concordance of this massively-parallel bisulphite sequencing with the microarray results for 2 of the 3 stably predicted DMRs, at the *OR2L13* and *FAM134B* genes, both showing decreased DNA methylation at these promoter loci. Of these, *OR2L13* has changes in DNA methylation that reaches statistical significance (p < 0.05) using a t test comparing the group of CGs tested by bisulphite sequencing in the region between the ASD and TD groups, with *FAM134B* also showing the decrease in DNA methylation predicted by the microarray study and trending towards but not reaching statistical significance (p = 0.0957). Interestingly, *OR2L13* has been found in two previous studies to demonstrate either altered DNA methylation in blood [9] or gene expression in brain [32] in individuals with ASD.

**Figure 3:**
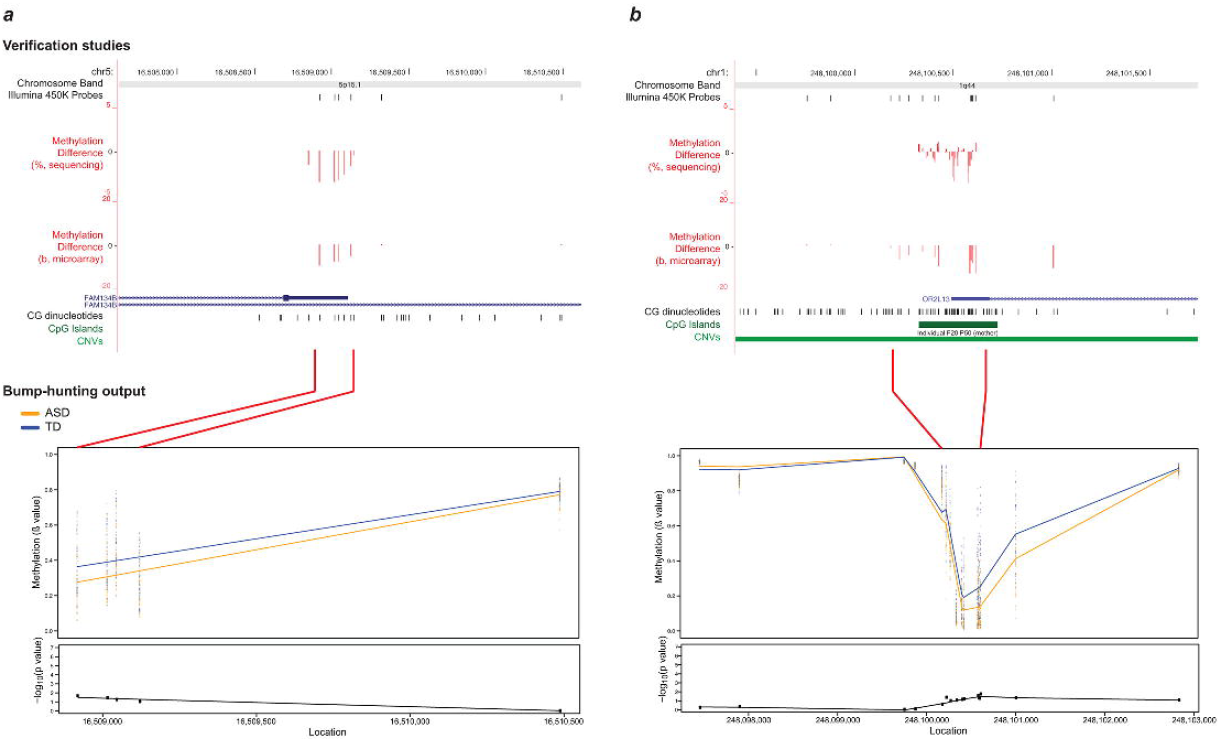
Massively-parallel bisulphite sequencing testing of candidate differentially methylated regions. Differences in DNA methylation between ASD and TD cohorts are shown for **(a)** *FAM134B* and **(b)** *OR2L13*. Absolute methylation values are displayed in the top panels, with the –log_10_ p-values as determined by bump-hunting (*dmrFind*). Differences in microarray mean ß value (ASD-TD) and massively-parallel bisulphite sequencing data (ASD-TD) show concordance for decreased DNA methylation in the ASD subjects at both loci (middle panels). The Illumina 450K Probes track displays CGs tiled by probes on the microarray. While the trends of DNA methylation changes were confirmed by the sequencing-based approaches, statistical significance testing was positive (p<0.05) for the *OR2L13* locus, with a trend towards significance at the *FAM134B* locus (split violin plots, lower panels). Of all the subjects tested, a CNV was found in only one individual at *OR2L13*, otherwise neither locus had CNVs present that could potentially affect interpretation of results.

We also explored the effect of the subjects’ age in more detail, identifying 306 potential DMRs (**Supplemental Table S6**) associated with the age of the individuals at the time of sample collection (which in our study ranged from 1-28 years, **Supplemental Table S1**). Gene ontology analysis demonstrated that the genes associated with these DMRs in buccal epithelium are significantly enriched in pathways especially related to development and differentiation but also include pathways involved in neuronal development (**Supplemental Figure S7**). It is notable that buccal epithelium should be informative for these patterns and supports the use of this cell type as a surrogate for studies of events occurring in the central nervous system.

### Co-methylation Network Analysis

While DMR identification is useful for the reasons described above, the exclusion from consideration of the majority of probes on the microarray in a bump-hunting approach and the failure of such approaches to consider that multiple dispersed loci may change DNA methylation patterns concordantly prompted our focus on a network-based analytical approach. Previous work has demonstrated the utility of employing such a network-based approach to detect expression and DNA methylation differences potentially missed by conventional statistical difference testing [32,33]. We therefore chose to use Weighted Gene Co-expression Network Analysis (WGCNA) [34] as a complement to bump-hunting, identifying instead of DMRs the patterns of co-methylation that distinguish the ASD subjects. We found co-methylation modules associated with all of our known biological covariates, including age, gender, and percentage of local ancestry. We selected modules significantly associated only with ASD case/control status after stringent Bonferroni correction. We obtained two modules that met these criteria; the full list of association p-values is provided in **Supplemental Table S7**. The larger co-methylation module significantly associated with ASD contained 116 CGs (Figure 4a). Genes in proximity to these CGs (**Supplemental Table S8**) include *NF1*, whose dysfunction is strongly associated with syndromic autism [35]. Ontology analysis of these genes revealed significant enrichment for functional categories including negative regulation of cell migration, locomotion, cellular compartment movement and cell proliferation, categories previously implicated in ASD subjects [2]. However, because of the recent recognition that gene ontology analysis can be biased towards genes with a greater number of probes [36], we used the number of probes per gene defined in the Illumina manifest to generate a weight for each gene, and recalculated enriched gene ontologies using the R package *GOseq* [37]. The p-values of the resulting candidate ontology categories did not survive correction for multiple testing, indicating that this kind of ontology analysis needs to be interpreted with caution, and may prompt reanalysis of prior, published gene ontological associations in ASD and other phenotypes.

**Figure 4:**
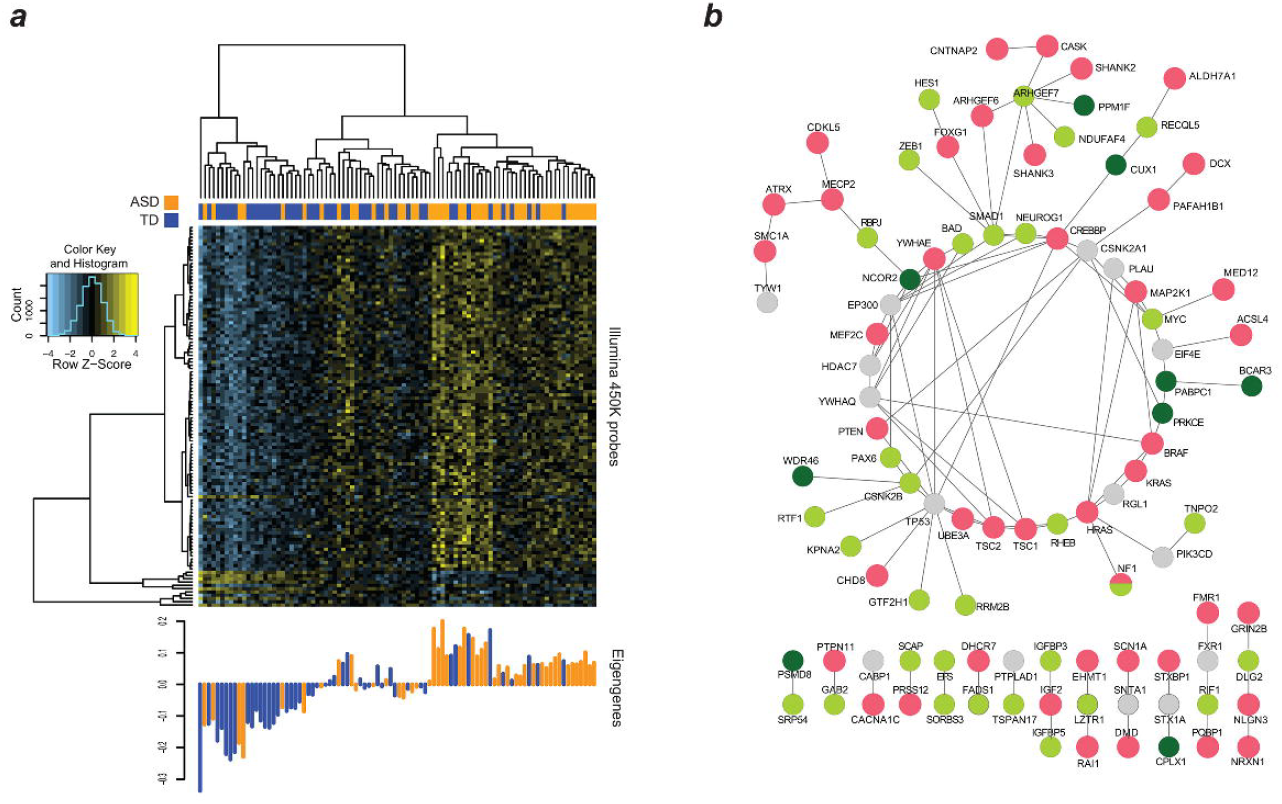
Methylation of CGs in WGCNA modules associated with ASD status. A. The heat map reflects unsupervised clustering of methylation values of CGs in the “light green” module significantly associated with ASD alone. Clear segregation of ASD (orange) and TD (blue) individuals can be seen in these CGs. The bottom panel shows the corresponding eigengenes for each individual.
B. Known ASD genes (red) and those from each of the two WGCNA modules (green shades) with connecting genes (grey) showing extensive interactions, and the linking of separated ASD gene groups by those identified in the current study.

As a different means of assessing the functional significance of the ASD-associated co-methylation modules, we tested whether they were functionally and non-randomly connected to known ASD risk genes. Using data from only the more stringent protein physical interaction databases [38], we created a protein-protein interaction network using the genes from the two significant WGCNA modules and a reference list of known ASD genes [39]. **Figure 4b** displays the resulting highly interconnected network, demonstrating that many of our module genes directly interact with genes previously found to be mutated in ASD. To test whether this degree of interaction was non-random, we performed the Degree-Aware Disease Gene Prioritization (DADA) approach [40] previously used to study associations of sporadic gene mutations with ASD [12]. We found that our module genes were significantly enriched in ranking (Mann Whitney U = 7.1 × 10^−3^), indicating the non-random selection of epigenetically-dysregulated genes for those within pathways previously implicated in ASD.

## DISCUSSION

Despite using a surrogate ectodermal cell type, this study revealed a pattern of altered DNA methylation at genes expressed in the brain, encoding proteins enriched for the post-natal synaptic transmission functions previously implicated in the pathogenesis of ASD, and significantly interactive with proteins encoded by genes previously described to be mutated in subjects with ASD. These findings combine with those of other recent studies to support a contribution of epigenetic dysregulation to the pathogenesis of ASD [8,9].

A current concern with the EWAS approach is that the generally small changes in DNA methylation found [17] may not be substantially in excess of the noise introduced by technical or biological effects influencing DNA methylation that have no relationship to the phenotype being tested. We have an increasing appreciation of the types of influences on DNA methylation in studies of human subjects, allowing us to design and execute studies more rigorously now than was appreciated to be necessary in the past. The current study represents the largest epigenome-wide analysis to date testing a single cell type in ASD. The use of such an homogeneous cell type in this study should minimize the problems recognized to be associated with samples of mixed cellularity [18]. Our parallel SNP genotyping allowed not only the detection and elimination of CNVs and mosaic aneuploidy as influences, it also allowed us to map within chromosomes the ancestral haplotype in which each site tested for DNA methylation was located, helping to control for genetic influences on DNA methylation. Analytically, we accounted for these sources of variability and used the SVA approach [41] from within *bump-hunting* [28] to identify and account for known influences (age [16] and sex [22]) as well as additional, otherwise unrecognized influences. The analysis also included iterative use of the data preprocessing and analysis steps prior to using *bump-hunting* to highlight the DMRs that are stably predicted, and gene ontology studies that address the newly-recognized concern that the number of microarray probes representing the gene can influence the outcome of analysis [36]. These measures, which were preceded by stringent pre-processing of the microarray data, appear to represent the currently necessary level of stringency for EWAS studies.

The partial changes in DNA methylation that are typical of EWAS imply an underlying property of the cell populations being studied. Unlike gene expression, which can vary quantitatively in an individual cell, DNA methylation is either present or absent on an allele, so that small changes of DNA methylation reflect allelic (presumably cellular) subpopulation mosaicism for the epigenetic changes. Our finding of limited changes in DNA methylation in buccal epithelium suggests a model for an early embryological event in ectodermal cells, perturbing the epigenome of a subpopulation of these cells, detectable postnatally in buccal epithelium but potentially also occurring as a mosaic cell subpopulation in other ectodermally-derived tissues including the brain. Mosaicism for chromosomal abnormalities has been associated with ASD [42-44], indicating that problems affecting only subsets of cells in the brain can lead to this condition, as also suggested in a recent review of somatic mosaicism in neurological disorders [45].

The characteristics of the genes at which the DNA methylation changes are occurring in the ASD cohort lend support to our model linking epigenetic disturbances with the pathogenesis of ASD. Despite studying non-neuronal cells, the differentially-methylated genes are those expressed in the brain, enriched for post-natal synaptic transmission function. Network approaches have proven to be an informative means of understanding the diversity of mutational events in ASD [32], prompting us to apply the same approaches to explore DNA methylation changes at the genes identified by WGCNA. We found that the genes in the most significant WGCNA modules are non-randomly enriched for interactions with genes already implicated in ASD. The model that results is of mosaic epigenetic dysregulation affecting the same networks and pathways targeted by mutational mechanisms, creating comparable deleterious effects on neuronal function. The results will need to be replicated to increase confidence in the conclusions, requiring independently-collected buccal epithelial samples from comparable subjects with ASD and suitable control individuals. As this currently represents an unusual source of clinical material, such a resource is not yet available, requiring that study replication will need to be addressed as the next phase of this project.

We conclude that of the two mechanisms we originally proposed for AMA causing ASD, covert aneuploidy occurring at detectable levels (≥20%) [46] is not as likely to be involved as epigenetic dysregulation. The use of mixed cell types in prior epigenetic studies of ASD [7] may have limited the ability to detect such subpopulation effects, although it is encouraging that, in a recent study of monozygotic twins, DNA methylation differences were also observed, implicating a gene that we also found to be targeted for epigenetic dysregulation, *OR2L13* [9]. This gene appears to be especially labile in ASD in terms of DNA methylation and expression – mixed leukocytes from the monozygotic twins study shows increased DNA methylation at this locus, whereas there is decreased DNA methylation in the ectodermal cells we studied, and expression of this gene is significantly upregulated in the Brodmann Area (BA) 44/45 and downregulated in the BA9/41 regions of the brain [32]. Whether this olfactory receptor gene is a contributor to the distinctive olfaction in subjects with ASD [47,48] remains to be determined, although it should also be noted that the Human Brain Transcriptome (HBT) studies categorized this gene, alone among all olfactory receptor genes, into a post-natal synaptic transmission co-expression module [31], indicating unique transcriptional properties for this gene among those encoding olfactory receptors. This gene merits further studies as being unusually prone to dysregulation in ASD.

Epigenetic changes observed in our cohort of subjects born to mothers with AMA may be due to aging of the oocyte, but there are additional potential mechanisms. Spermatozoa from the generally older fathers of the ASD subjects may also have epigenetic changes with aging. Epigenetic dysregulation in ASD could be due to as-yet unrecognized environmental influences during development. Furthermore, an emerging property of the genes in which mutations or sequence variants occur in subjects with ASD is that of chromatin and epigenetic regulatory properties [11], which may be occurring in ASD subjects born to older mothers as a consequence of the generally older paternal age of such individuals and the increased risk of mutational events with age in the male germline [12,13]. The epigenetic dysregulation we observe may therefore represent a common outcome of the effects of mutations in such genes, making epigenetic changes an event secondary to genetic mutations within cohorts of subjects with ASD. Combined genetic and epigenetic analyses of the same subjects will be needed to test these possibilities.

## MATERIALS & METHODS

### Cohorts and sample collection

All patient recruitment and sample collection was performed with the appropriate human subjects protocol approval from the Institutional Review Board at the Albert Einstein College of Medicine. We enrolled two groups of subjects: individuals with a diagnosed autism spectrum disorder (ASD), born to mothers aged 35 and older, and a control group of typically developing (TD) individuals born to mothers aged 35 and older. Of these, we selected 50 ASD and 50 TD subjects for genotyping analysis, of which we took 47 and 48 from each group, respectively, for methylation analysis. The characteristics of the ASD and TD cohorts are summarized in **Supplemental Table S1**.

We collected buccal epithelium using exfoliative brushing, choosing this accessible cell type for two major reasons. Firstly, as ectodermal in origin, these cells have been found to represent a better surrogate for the brain than other tissues when relating cognitive impairment with tissue-specific levels of mosaic trisomy 21 [49]. Secondly, exfoliative buccal brushing harvests squamous epithelial cells homogeneously, confirmed by microscopy of samples in this study (**Supplemental Figure S1**), avoiding the potentially confounding effects of heterogeneous cell subtypes on epigenomic profiling [18]. DNA from the buccal epithelial samples was extracted with a modification of the protocol for the Qiagen Gentra Puregene Buccal Cell Kit. We randomized samples between arrays as a precaution against batch effects and ran them on the Illumina HumanOmni2.5-8 BeadChip genotyping platform and the Illumina Infinium HumanMethylation450 Beadchip methylation platform (**Supplemental Table S2**).

### Chromosomal mosaicism analysis

Stringent genotype data preprocessing included filtering for poor quality arrays and probes, as well as probe missingness and allele frequency (**Text S1**). After this preprocessing, we used the Mosaic Alteration Detection (*MAD*) algorithm [27] to assess the prevalence of chromosomal mosaicism. We confirmed the ability of MAD to detect mosaicism in samples from 4 individuals with known chromosomal mosaicism (three cell line and one buccal epithelial samples, **Supplemental Figure S2**).

### Local ancestry deconvolution

We used the *HAPMIX* [50] software to annotate genomic ancestry genome-wide. Since our population comprised individuals with 3-way admixture, we simulated mixed parental ancestry and calculated probabilities of each potential genotype at every probe. We assigned ancestry only when the local probability exceeded 0.5 (**Supplemental Table S3**).

### Differential methylation analysis

Arrays were subjected to stringent preprocessing to correct for technical (probe type, detection p-value), and batch artefacts (**Supplemental Figure S3**). Due to inclusion of both males and females in our cohort, we removed the sex chromosomes from our dataset and conducted analysis on autosomes only. Detailed information on the pre-processing procedures is included in the **Supplementary Information**.

After preprocessing, we then performed principal components analysis (PCA) on the M values (logit-transformed Illumina-defined beta values) obtained. We accounted for the possible known confounding influences, including technical (date of DNA extraction, microarray chip, position on chip), microarray-based (all categories of control probes designed by Illumina) and biological (ASD status, age, gender, and ancestry percentage). Ancestry percent was calculated as the proportion for each population of all allele genotyping positions called by *HAPMIX*. We identified the significant confounding covariates and corrected for them prior to all subsequent analysis.

To identify differentially methylated regions (DMRs), we used the bump-hunting approach, specifically the *dmrFind* algorithm within the *charm* package [28]. Based on our PCA data, we input age, gender, percent CEU (European) and percent YRI (African) ancestry as known biological covariates. Those loci called as significant by the *dmrFind* algorithm were defined as candidate DMRs. Stability of DMR prediction was tested by re-running the bump-hunting on the data a total of 4 times. Additional loci identified through bump-hunting are illustrated in **Supplemental Figure S5**.

### Copy number variant (CNV) calling

We utilized the *CNVision* algorithm [30] to detect copy number variants in all individuals. We removed samples with CNVs overlapping called DMRs and re-ran bump-hunting to confirm that the DMR was not solely due to the presence of CNVs in the region.

### Weighted gene co-expression network analysis (WGCNA)

We used the Weighted Correlation Network Analysis framework *WGCNA* [34] for our DNA methylation analysis, as has previously been performed [33] to identify networks of co-methylated CG dinucleotides associated with ASD status. As input, we used the CG values generated from the Surrogate Variable Analysis (*SVA*) algorithm called in the *dmrFind* function performing bump-hunting, thus correcting for all known technical artifacts. We used the *WGCNA* package in R and built an unsigned co-methylation network. Correlation matrices were raised to the power of 5, as calculated by the scale-free topology criterion on data subsets, and thresholds were set of minimum module connectivity (kME) of greater than 0.7, and minimum height for module merging of 0.1 [32]. We ran the algorithm with block sizes of 40,000 CGs.

We assessed module relevance to case/control status with a t-test (two tailed, unequal variance) of module eigengene values with case/control and gender categories. For relationship with the continuous variables of age and percent of YRI and CEU ancestry, we used Pearson correlation coefficients and their Student asymptotic p-values.

For analysis of methylation changes associated with ASD, we selected the 2 modules (“light green”, and “dark olive green2”) that showed significant correlation only with ASD status and not with any other covariate, to avoid introducing confounding effects. The genes associated with the dark olive green2 module are listed in **Supplemental Table S9**.

### Functional Analysis

The nature of the enriched pathways involved in the list of genes associated with DMRs was investigated using a Cytoscape plugin for Biological Network Gene Ontology Enrichment, *BiNGO* [51]. The reference set used was set as the whole annotation from *Homo sapiens* and Biological Processes from Gene Ontology were queried, with a significance level set to a false discovery rate (FDR) corrected value of 0.05.

To assess the functional impact of our WGCNA ASD-associated co-methylated gene modules, we interrogated the genes’ relevance in protein-protein interaction (PPI) networks. We combined the genes in the light green and dark olive green 2 modules with a list of previously curated known ASD risk genes, with the addition of exome sequencing candidates (*KATNAL2* and *CHD8*) [39]. We used *GeneMania* [38] to build a PPI network of this combined list, using only data from physical protein interaction databases. We visualized this network in Cytoscape. To test for bias due to probe number differences at each gene, we use the *GoSeq* package written in R [37] with weighting of genes defined by the number of probes assigned to each gene in the Illumina 450K microarray manifest. A PPI analysis showing interactions with intellectual disability (ID) genes is shown in **Supplemental Figure S8**.

To test the significance of these PPI connections, we performed Degree Aware Disease Gene Prioritization (*DADA*) [40]. We used the ASD seed list mentioned previously, a combined candidate list of the genes related to the ASD-associated WGCNA CGs, and the physical interaction database from the Human Protein Reference Database (HPRD) available through *GeneMania*. We evaluated if our genes were significantly enriched in ranking using the Mann-Whitney test.

### Verification of differential methylation

We bisulphite converted 500 ng of DNA using the Zymo EZ-96 Methylation-Lightning Kit. After separate PCR amplification of target regions (primers shown in **Supplemental Table S5**), we pooled the amplicons in averaged equal ratios and generated Illumina libraries using Tecan automation. Two sets of 48 libraries each were multiplexed on the MiSeq. Using *bsmap* (Bisulphite Sequencing Mapping Platform) [52] we checked for bisulphite conversion efficiency (C→T in CH contexts) and quantified the percent methylation for each person at every CG in the amplicons. Additional loci tested by this bisulphite sequencing approach are shown in **Supplemental Figure S6**.

### Data access

All microarray data generated are deposited into the Gene Expression Omnibus (GEO) database under accession number GSE50759.

## ACKNOWLEDGMENTS

Autism Speaks is thanked for helping with patient recruitment.

## SUPPLEMENTARY DATA

### Text S1

A file containing all supplementary figures and tables, with details on analytical approaches and including software allowing re-analysis of data.

## REFERENCES

1. Morrow EM (2010) Genomic copy number variation in disorders of cognitive development. J Am Acad Child Adolesc Psychiatry 49: 1091–1104.

2. Pinto D, Pagnamenta AT, Klei L, Anney R, Merico D, et al. (2010) Functional impact of global rare copy number variation in autism spectrum disorders. Nature 466: 368–372.

3. Muers M (2012) Human genetics: Fruits of exome sequencing for autism. Nat Rev Genet 13: 377.

4. Devlin B, Scherer SW (2012) Genetic architecture in autism spectrum disorder. Curr Opin Genet Dev 22: 229–237.

5. Grabrucker AM (2012) Environmental factors in autism. Front Psychiatry 3: 118.

6. Miyake K, Hirasawa T, Koide T, Kubota T (2012) Epigenetics in autism and other neurodevelopmental diseases. Adv Exp Med Biol 724: 91–98.

7. Ginsberg MR, Rubin RA, Falcone T, Ting AH, Natowicz MR (2012) Brain transcriptional and epigenetic associations with autism. PLoS One 7: e44736.

8. Shulha HP, Cheung I, Whittle C, Wang J, Virgil D, et al. (2012) Epigenetic signatures of autism: trimethylated H3K4 landscapes in prefrontal neurons. Arch Gen Psychiatry 69: 314–324.

9. Wong CC, Meaburn EL, Ronald A, Price TS, Jeffries AR, et al. (2013) Methylomic analysis of monozygotic twins discordant for autism spectrum disorder and related behavioural traits. Mol Psychiatry.

10. Ladd-Acosta C, Hansen KD, Briem E, Fallin MD, Kaufmann WE, et al. (2013) Common DNA methylation alterations in multiple brain regions in autism. Mol Psychiatry.

11. Ben-David E, Shifman S (2013) Combined analysis of exome sequencing points toward a major role for transcription regulation during brain development in autism. Mol Psychiatry 18: 1054–1056.

12. O’Roak BJ, Vives L, Girirajan S, Karakoc E, Krumm N, et al. (2012) Sporadic autism exomes reveal a highly interconnected protein network of de novo mutations. Nature 485: 246–250.

13. Kong A, Frigge ML, Masson G, Besenbacher S, Sulem P, et al. (2012) Rate of de novo mutations and the importance of father’s age to disease risk. Nature 488: 471–475.

14. Sandin S, Hultman CM, Kolevzon A, Gross R, MacCabe JH, et al. (2012) Advancing maternal age is associated with increasing risk for autism: a review and meta-analysis. J Am Acad Child Adolesc Psychiatry 51: 477–486 e471.

15. Pellestor F, Anahory T, Hamamah S (2005) Effect of maternal age on the frequency of cytogenetic abnormalities in human oocytes. Cytogenet Genome Res 111: 206–212.

16. Heyn H, Li N, Ferreira HJ, Moran S, Pisano DG, et al. (2012) Distinct DNA methylomes of newborns and centenarians. Proc Natl Acad Sci U S A 109: 10522–10527.

17. Rakyan VK, Down TA, Balding DJ, Beck S (2011) Epigenome-wide association studies for common human diseases. Nat Rev Genet 12: 529–541.

18. Houseman EA, Accomando WP, Koestler DC, Christensen BC, Marsit CJ, et al. (2012) DNA methylation arrays as surrogate measures of cell mixture distribution. BMC Bioinformatics 13: 86.

19. Wang D, Yan L, Hu Q, Sucheston LE, Higgins MJ, et al. (2012) IMA: an R package for high-throughput analysis of Illumina’s 450K Infinium methylation data. Bioinformatics 28: 729–730.

20. Boks MP, Derks EM, Weisenberger DJ, Strengman E, Janson E, et al. (2009) The relationship of DNA methylation with age, gender and genotype in twins and healthy controls. PLoS One 4: e6767.

21. Bell JT, Tsai PC, Yang TP, Pidsley R, Nisbet J, et al. (2012) Epigenome-wide scans identify differentially methylated regions for age and age-related phenotypes in a healthy ageing population. PLoS Genet 8: e1002629.

22. Sarter B, Long TI, Tsong WH, Koh WP, Yu MC, et al. (2005) Sex differential in methylation patterns of selected genes in Singapore Chinese. Hum Genet 117: 402–403.

23. Bell JT, Pai AA, Pickrell JK, Gaffney DJ, Pique-Regi R, et al. (2011) DNA methylation patterns associate with genetic and gene expression variation in HapMap cell lines. Genome Biol 12: R10.

24. Gibbs JR, van der Brug MP, Hernandez DG, Traynor BJ, Nalls MA, et al. (2010) Abundant quantitative trait loci exist for DNA methylation and gene expression in human brain. PLoS Genet 6: e1000952.

25. Gertz J, Varley KE, Reddy TE, Bowling KM, Pauli F, et al. (2011) Analysis of DNA methylation in a three-generation family reveals widespread genetic influence on epigenetic regulation. PLoS Genet 7: e1002228.

26. Teschendorff AE, Menon U, Gentry-Maharaj A, Ramus SJ, Gayther SA, et al. (2009) An epigenetic signature in peripheral blood predicts active ovarian cancer. PLoS One 4: e8274.

27. Gonzalez JR, Rodriguez-Santiago B, Caceres A, Pique-Regi R, Rothman N, et al. (2011) A fast and accurate method to detect allelic genomic imbalances underlying mosaic rearrangements using SNP array data. BMC Bioinformatics 12: 166.

28. Jaffe AE, Murakami P, Lee H, Leek JT, Fallin MD, et al. (2012) Bump hunting to identify differentially methylated regions in epigenetic epidemiology studies. Int J Epidemiol 41: 200–209.

29. Chen YA, Lemire M, Choufani S, Butcher DT, Grafodatskaya D, et al. (2013) Discovery of cross-reactive probes and polymorphic CpGs in the Illumina Infinium HumanMethylation450 microarray. Epigenetics 8: 203–209.

30. Sanders SJ, Ercan-Sencicek AG, Hus V, Luo R, Murtha MT, et al. (2011) Multiple recurrent de novo CNVs, including duplications of the 7q11.23 Williams syndrome region, are strongly associated with autism. Neuron 70: 863–885.

31. Kang HJ, Kawasawa YI, Cheng F, Zhu Y, Xu X, et al. (2011) Spatio-temporal transcriptome of the human brain. Nature 478: 483–489.

32. Voineagu I, Wang X, Johnston P, Lowe JK, Tian Y, et al. (2011) Transcriptomic analysis of autistic brain reveals convergent molecular pathology. Nature 474: 380–384.

33. van Eijk KR, de Jong S, Boks MP, Langeveld T, Colas F, et al. (2012) Genetic analysis of DNA methylation and gene expression levels in whole blood of healthy human subjects. BMC Genomics 13: 636.

34. Langfelder P, Horvath S (2008) WGCNA: an R package for weighted correlation network analysis. BMC Bioinformatics 9: 559.

35. Garg S, Lehtonen A, Huson SM, Emsley R, Trump D, et al. (2013) Autism and other psychiatric comorbidity in neurofibromatosis type 1: evidence from a population-based study. Dev Med Child Neurol 55: 139–145.

36. Geeleher P, Hartnett L, Egan LJ, Golden A, Raja Ali RA, et al. (2013) Gene-set analysis is severely biased when applied to genome-wide methylation data. Bioinformatics 29: 1851–1857.

37. Young MD, Wakefield MJ, Smyth GK, Oshlack A (2010) Gene ontology analysis for RNA-seq: accounting for selection bias. Genome Biol 11: R14.

38. Warde-Farley D, Donaldson SL, Comes O, Zuberi K, Badrawi R, et al. (2010) The GeneMANIA prediction server: biological network integration for gene prioritization and predicting gene function. Nucleic Acids Res 38: W214–220.

39. Neale BM, Kou Y, Liu L, Ma’ayan A, Samocha KE, et al. (2012) Patterns and rates of exonic de novo mutations in autism spectrum disorders. Nature 485: 242–245.

40. Erten S, Bebek G, Ewing RM, Koyuturk M (2011) DADA: Degree-Aware Algorithms for Network-Based Disease Gene Prioritization. BioData Min 4: 19.

41. Leek JT, Johnson WE, Parker HS, Jaffe AE, Storey JD (2012) The sva package for removing batch effects and other unwanted variation in high-throughput experiments. Bioinformatics 28: 882–883.

42. Abu-Amero KK, Hellani AM, Salih MA, Seidahmed MZ, Elmalik TS, et al. (2010) A de novo marker chromosome derived from 9p in a patient with 9p partial duplication syndrome and autism features: genotype-phenotype correlation. BMC Med Genet 11: 135.

43. Chen CP, Lin SP, Su JW, Lee MS, Wang W (2012) Phenotypic features associated with mosaic tetrasomy 9p in a 20-year-old female patient include autism spectrum disorder. Genet Couns 23: 335–338.

44. Kostanecka A, Close LB, Izumi K, Krantz ID, Pipan M (2012) Developmental and behavioral characteristics of individuals with Pallister-Killian syndrome. Am J Med Genet A 158A: 3018–3025.

45. Poduri A, Evrony GD, Cai X, Walsh CA (2013) Somatic mutation, genomic variation, and neurological disease. Science 341: 1237758.

46. Jacobs KB, Yeager M, Zhou W, Wacholder S, Wang Z, et al. (2012) Detectable clonal mosaicism and its relationship to aging and cancer. Nat Genet 44: 651–658.

47. Bennetto L, Kuschner ES, Hyman SL (2007) Olfaction and taste processing in autism. Biol Psychiatry 62: 1015–1021.

48. Galle SA, Courchesne V, Mottron L, Frasnelli J (2013) Olfaction in the autism spectrum. Perception 42: 341–355.

49. Papavassiliou P, York TP, Gursoy N, Hill G, Nicely LV, et al. (2009) The phenotype of persons having mosaicism for trisomy 21/Down syndrome reflects the percentage of trisomic cells present in different tissues. Am J Med Genet A 149A: 573–583.

50. Price AL, Tandon A, Patterson N, Barnes KC, Rafaels N, et al. (2009) Sensitive detection of chromosomal segments of distinct ancestry in admixed populations. PLoS Genet 5: e1000519.

51. Maere S, Heymans K, Kuiper M (2005) BiNGO: a Cytoscape plugin to assess overrepresentation of gene ontology categories in biological networks. Bioinformatics 21: 3448–3449.

52. Xi Y, Li W (2009) BSMAP: whole genome bisulfite sequence MAPping program. BMC Bioinformatics 10: 232.

53. Risch N, Spiker D, Lotspeich L, Nouri N, Hinds D, et al. (1999) A genomic screen of autism: evidence for a multilocus etiology. Am J Hum Genet 65: 493–507.

54. Myers RA, Casals F, Gauthier J, Hamdan FF, Keebler J, et al. (2011) A population genetic approach to mapping neurological disorder genes using deep resequencing. PLoS Genet 7: e1001318.

55. Lauritsen MB, Als TD, Dahl HA, Flint TJ, Wang AG, et al. (2006) A genome-wide search for alleles and haplotypes associated with autism and related pervasive developmental disorders on the Faroe Islands. Mol Psychiatry 11: 37–46.

56. Nishimura Y, Martin CL, Vazquez-Lopez A, Spence SJ, Alvarez-Retuerto AI, et al. (2007) Genome-wide expression profiling of lymphoblastoid cell lines distinguishes different forms of autism and reveals shared pathways. Hum Mol Genet 16: 1682–1698.

